# Drug Repurposing of Rapamycin Mitigates Amyloid-β-Associated Proteotoxicity in Caenorhabditis elegans Models of Alzheimer’s Disease

**DOI:** 10.64898/2026.11.14.766704

**Authors:** Samirah Alkhudaydi, Abdullatif Alsulami, Makayla Linfoot, Megan Terlouw, Ava Holzer, Faisal Albalawi, Stephanie McGrath, Julie A. Moreno

## Abstract

**Background:** Alzheimer’s disease (AD) is the leading cause of dementia worldwide, affecting an estimated 55 million people and projected to exceed 130 million by 2050. AD is defined pathologically by extracellular accumulation of amyloid-beta (Aβ) and intracellular neurofibrillary tangles. Currently no therapy halts the underlying neurodegeneration in AD patients. Proteotoxic accumulation of Aβ is a central pathogenic event therefore, strategies that enhance its clearance represent a compelling therapeutic avenue. Rapamycin (sirolimus), an FDA-approved allosteric inhibitor of the mechanistic target of rapamycin complex 1 (mTORC1) that relieves mTORC1-mediated suppression of autophagy, positioning it as an attractive drug-repurposing candidate for AD. However, it remains unclear whether rapamycin’s neuroprotection depends on directly reducing Aβ burden or acts through Aβ- independent mechanisms. This question, along with whether rapamycin alters specific Aβ species, has not been systematically examined in a single, genetically tractable model.

**Methods/Results:** Using two transgenic *C. elegans* models of human Aβ_1–42_ expression a temperature-inducible, muscle-specific strain CL4176 and the pan-neuronal strain CL2355. We exposed *C. elegans* to 20 μM rapamycin the lowest, non-toxic concentration to produce a protective effect. Rapamycin treatment delayed Aβ-induced paralysis in log-rank p = 0.0001) and improved crawling and swimming locomotion parameters. In the pan-neuronal CL2355 strain, rapamycin rescued the Aβ-induced chemotaxis deficit (one-way ANOVA, p = 0.008) without altering chemotaxis in the non-transgenic CL2122 control. Rapamycin also extended Kaplan–Meier survival in CL2355 (log-rank, Bonferroni-adjusted p < 2 × 10⁻¹⁶) and preserved locomotor health span, delaying the trajectory of age-related motor decline. At the molecular level, rapamycin did not significantly affect Aβ burden.

**Conclusion:** Rapamycin provides broad-spectrum and Aβ-selective neuroprotection across muscle-and neuron-expressing *C. elegans* models of Aβ proteotoxicity, extending from paralysis and locomotion to chemosensory behavior, longevity, and health span.

## Introduction

Alzheimer’s disease (AD) is a chronic, age-associated neurodegenerative disorder and the most common cause of dementia, accounting for 60–70% of cases and currently affecting an estimated 55 million people worldwide ^1^, with prevalence projected to exceed 130 million by 2050 ^2^. AD is defined pathologically by two hallmark lesions: extracellular accumulation of amyloid-beta (Aβ) peptides and intracellular neurofibrillary tangles composed of hyperphosphorylated tau ^3^. Aβ is generated by sequential proteolytic cleavage of the amyloid precursor protein (APP) by β- and γ-secretases; the resulting Aβ_1–42_ isoform is more aggregation-prone than Aβ1–40 species ^4,5^. The imbalance between production and clearance of the proteins is the core of the amyloid cascade hypothesis that Aβ accumulation initiates a cascade of tau pathology, neuroinflammation, and neurodegeneration ^4,6^.

Aging is the dominant risk factor for late-onset AD ^2^, with several cellular hallmarks including: loss of proteostasis, mitochondrial dysfunction and disabled autophagy which will contribute to AD pathogenesis by reducing neuronal capacity to clear misfolded Aβ and tau species ^6,7^. Rapamycin (sirolimus), an FDA-approved drug that allosterically inhibits mTORC1 by binding FKBP12, enhances the autophagy that is known to be elevated in post-mortem AD brain and in transgenic AD mouse models ^8,9^. It is also one of the most robust lifespan-extending pharmacological agents identified to date, extending lifespan across yeast, *C. elegans*, Drosophila, and mice ^9^. In transgenic rodent AD models, rapamycin reduces Aβ plaque burden, enhances autophagic flux and reduces neuroinflammation ^10,11^.

Conventional drug discovery takes 15–20 years and over $1 billion per approved drug.². This pace poorly suited to a disease whose prevalence is expected to more than double within 30 years.² Drug repurposing redirects approved compounds with established safety and pharmacokinetic profiles to new indications, offering a faster, cheaper and lower-risk alternative¹².Progress in AD drug discovery remains constrained by the cost and throughput limitations of mammalian models ^12^. The nematode C.elegans offers a genetically tractable, rapidly scalable alternative, with a short lifespan, fully mapped nervous system, and conserved proteostatic machinery amenable to high-throughput pharmacology ^13^. Transgenic strains engineered to express human Aβ_1–42_ in specific tissues allow proteotoxicity to be isolated from upstream APP processing^14^: CL4176 (*smg-1(cc546)* I; *dvIs27* X) expresses Aβ_1–42_ in body-wall muscle under a temperature-inducible promoter, producing a quantitative, pharmacologically sensitive paralysis phenotype^14,15^, while CL2355 (*smg-1(cc546) dvIs50* I) expresses Aβ_1–42_ pan-neuronally, producing behavioral deficits more directly analogous to the cognitive and functional phenotypes of human AD ^16,17^.

Although rapamycin is neuroprotective in mammalian AD models ^10^, it is unknown how rapamycin dependent reduction in Aβ proteotoxicity effects motor, chemosensory, survival, and healthy-aging outcomes. To address this question, we have utilized muscle-and neuron-Aβ expressing *C. elegans* models. We found that rapamycin selectively protects against Aβ proteotoxicity in both muscle-and neuron-expressing *C. elegans* models, improving paralysis, locomotion, chemosensory, lifespan and healthspan.

## Materials and Methods

### *C. elegans* strains, maintenance, and synchronization

Four Caenorhabditis elegans strains were obtained from the Caenorhabditis Genetics Center (University of Minnesota, Minneapolis, MN, USA): CL4176 (dvIs27 [myo-3p::Abeta(1-42)::let-851 3’UTR) + rol-6(su1006)]), which expresses human Aβ_1–42_ in body-wall muscle under a temperature-inducible myo-3 promoter and expression via smg-1 ^14,15^; CL2355 (dvIs50[Psnb-1:: Aβ_1–42_::3′UTR(long) + Pmtl-2::GFP]), which drives pan-neuronal Aβ_1–42_ expression following temperature upshift ^17^; and the genetically matched non-Aβ control strains CL802 (smg-1(cc546) I; rol-6(su1006) II) and CL2122 (dvIs15 [(pPD30.38) unc-54(vector) + (pCL26) mtl-2::GFP]). Worms were maintained on nematode growth medium (NGM) seeded with E. coli OP50 at 16 °C _18_. Transgene expression was induced experimentally by upshifting to 25 °C ^13^. Age-synchronized populations were obtained by alkaline hypochlorite (bleaching) treatment of gravid adults, followed by overnight hatching of released eggs on unseeded NGM at 16 °C to arrest larvae at the L1 stage ^18,19^.

### Rapamycin preparation

Rapamycin (sirolimus; Cayman Chemical, Ann Arbor, MI, USA; ≥98% purity) was dissolved in dimethyl sulfoxide (DMSO) to a 10 mM. Rapamycin stock is diluted into molten NGM that is chilled at 40°C before pouring; final DMSO concentration never exceeded 0.2% (v/v) in experimental plates. Vehicle treatment is 0.2% DMSO (Sigma-Aldrich, 47230).

### Amyloid-β-induced paralysis assay

The temperature-inducible paralysis assay was performed in CL4176 following Dostal and Link _14_. Synchronized L1 larvae were placed on NGM containing 0.2% DMSO (vehicle) or 20μM rapamycin, seeded with OP50, and cultured at 16 °C for 36 h before upshifting to 25 °C to induce muscle-specific Aβ_1–42_ expression (designated 0 h). Beginning at 20 h post-induction, paralysis was scored every 2 h until all vehicle-treated worms were paralyzed; a worm was scored as paralyzed if it failed to propagate a full sinusoidal body wave or moved only its head region in response to gentle prodding with a platinum wire pick ^20^. Each condition included ≥50 worms. The time at which 50% of the population was paralyzed (PT50) was calculated by linear interpolation between the two bracketing time points.

### Crawling and swimming locomotion analysis

Crawling and swimming locomotion assays were carried out using the body-wall muscle Aβ_1–42_ expressing strain (CL4176) and the non Aβ control strain (CL802). Age-synchronized L1 were grown on Rapamycin or vehicle treated NGM plates for 36h at 16°C, then temperature was upshifted for another 36h at 25°C. The day of the assay worm’s density in M9 buffer was determined, and a drop of the worm suspension places on unseeded NGM and video-recorded for 1 min using a Zeiss microscope equipped with a recording camera for swimming tacking. Plates left to air dry to record the worms crawling. Videos are analyzed with WormLab imaging software (MBF Bioscience version 2026). Morphology analysis of the worm development was determined using Mean Worm Length (uM) and Mean Width (uM). For crawling we assessed Self-Contact Distance (curling), number of reversals and straight-line distance. Similarly, swimming locomotion was assessed in M9 buffer, quantifying wave initiation rate, dynamic amplitude, and body wave number. All locomotion analysis was extracted per worm.

### Chemotaxis assay

Chemotaxis was assessed in the pan-neuronal muscle Aβ_1–42_ expressing strain (CL2355) and the non-Aβ transgenic strain (CL2122) following Bargmann et al. and Margie et al. ^21,22^ with some modifications. Briefly, age-synchronized worms were conditioned with either a vehicle or 20 μM rapamycin for 36 h at 16 °C, then the temperature was upshifted to 25 °C for 36 h to induce neuronal Aβ_1–42_ expression. Approximately 65 worms were placed at the center of a chemotaxis plate marked with two attractant spots (0.1% benzaldehyde) and two control spots (100% ethanol), each co-spotted with 0.25 M sodium azide to immobilize arriving worms. After 1 h at room temperature, the chemotaxis index (CI) was calculated as CI = (N attractant – N control)/N total.

### Lifespan and combined survival-locomotion (healthspan) assay

Lifespan was assessed in CL2355 and CL2122 following liquid media assay from Amrit FRG ^23^. Synchronized L4 larvae (∼100 worms/condition, distributed across 24-well plates) were maintained in medium containing 100 μM 5-fluoro-2′-deoxyuridine (FUDR) to suppress progeny production ^24^, with vehicle or 20 μM rapamycin. Transgene expression was induced by upshifting to 25 °C; 24 h post-upshift was designated day 1 of adulthood. Survival was scored every 48 h by response to gentle plate tabbing stimulation and pharyngeal pumping, log-rank (Mantel–Cox) tests with Bonferroni correction compared to survival curves. In the same cohort, locomotor activity (overall activity index) was video-recorded every time worms scored for lifespan and analyzed by WormLab to assess healthspan to investigate the locomotor behavior in pan-neuronally Aβ-expressing *C. elegans*.

### Western blotting

Worms were cultured with vehicle or 20 μM rapamycin for 36 h at 16 °C, then upshifted to 25 °C for 36 h. Worms density were determined and washed with miliqH2O and lysed by mixing with RIPA lysis buffer and freeze-thawed 3 times before sonication to rupture the worm’s cuticles then centrifuged 15000g for 15min at 4 °C. Total protein in the lysate was determined using BCA assay (Thermo Scintific, VI313751). 15 μg of total protein were resolved by SDS-PAGE (16.5% Tris-tricine gels for monomeric and oligomeric Aβ species) and transferred to 0.22 nitrocellulose membranes (Azure biosystems, 260406-09).^25^ Aβ was detected using the 1:500 6E10 mouse monoclonal antibody (BioLegend, San Diego, CA, USA; catalogue no. 803014) in 5% milk TBS-T blocking buffer and Horseradish peroxidase (HRP)-conjugated goat anti-mouse secondary antibody was obtained from Bio-Rad (catalog no.1706516). β-actin at 1:10000 used as a loading control from ABclonal (Woburn, MA, USA; catalog no.AC004). Band intensities were quantified in ImageJ/Fiji (v. 2.16.0/1.54p) ^26^ and normalized to β-actin.

## Statistical analysis

Data are presented as mean ± SEM. Analyses were performed in GraphPad Prism v11.0.1 (GraphPad Software) and R (v4.4.0+). Multiple-group comparisons used one-way ANOVA or for non-parametric data, the Kruskal–Wallis test. Two-group comparisons used unpaired two-tailed Student’s t-tests. Paralysis and survival data were analyzed by the Kaplan–Meier method with log-rank (Mantel–Cox) tests. Outliers were identified using ROUT method (Q=1%) in GraphPad Prism and excluded from analysis. All tests were done in triplicates unless otherwise noted. Significance was set at p < 0.05 (*p < 0.05, **p < 0.01, ***p < 0.001, ****p < 0.0001).

## Results

### Rapamycin delays Aβ-induced paralysis in CL4176

At 20μM, rapamycin significantly delayed the onset and progression of Aβ-induced paralysis relative to vehicle across the full observation window (log-rank, Bonferroni-adjusted p = 0.0001; Figure 1A), with no paralysis phenotype observed in the non-transgenic control strain CL802. The time to 50% population paralysis (PT50) post temperature up-shift showed no significance (Figure 1B). Paralysis phenotype was shown in the representative images (Figure 1C).

**Fig. 1.**
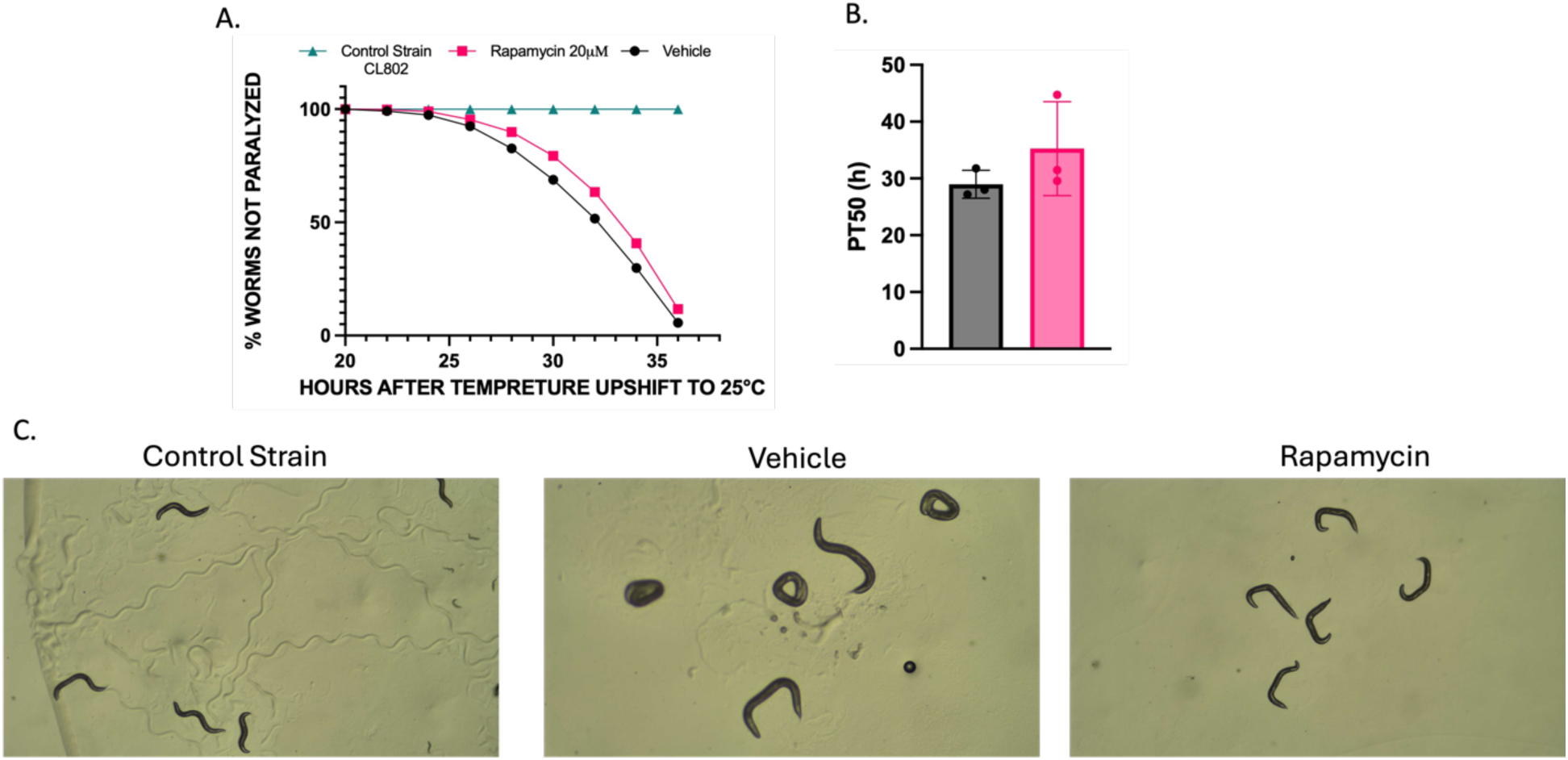
Rapamycin delays Amyloid Beta induced paralysis in muscle strain. **(A)** Paralysis assay CL4176 (Aβ-expressing) worms treated with vehicle (black) or 20 µM rapamycin (magenta), and the non-transgenic control strain CL802 (green), were scored for paralysis every 2 hours following temperature upshift to 25°C to induce Aβ expression. Significance is calculated by log-rank (Mantel–Cox) test (p < 0.0001, Bonferroni-corrected), n = 3 biological replicates per condition (∼150 worms per condition). **(B)** PT50 is not significant (time to 50% paralysis), calculated for each biological replicate by linear interpolation between the two time points flanking 50% paralysis (PT50 = t1 + [(50−p1)/(p2−p1)]×(t2−t1)). Bars show mean ± SED; points show the PT50 for each of the 3 individual biological replicates. **(C)** Representative images of CL4176 worms and control strain at the time of scoring.

### Rapamycin alters C.elegans development and enhances crawling and swimming locomotion in CL4176 strain

To determine the animal’s development, we examined worms that were treated with rapamycin or vehicle using WormLab software. Locomotion parameters (Table 1) used to quantify crawling and swimming behavior following Aβ induction. Body morphology showed that rapamycin-treated group demonstrates lower worm length but higher width in comparison with vehicle group (Figure 2A-B) On solid media, rapamycin-treated worms showed no significant difference from vehicle in self-contact distance (Figure 2C). On the other hand, rapamycin shows higher number of reversals from the vehicle group but remains insignificant from the control group (Figure 2D). Similar results are shown in the straight-line distance where rapamycin shows higher number of reversals from the vehicle group but remains insignificant from the control group (Figure 2E). In M9 buffer, rapamycin-treated worms showes significantly increased wave initiation rate (p <0.001), dynamic amplitude (p 0.0476), together with increased body wave number (p < 0.0065) (Figure 2, F-H).

**Figure 2.**
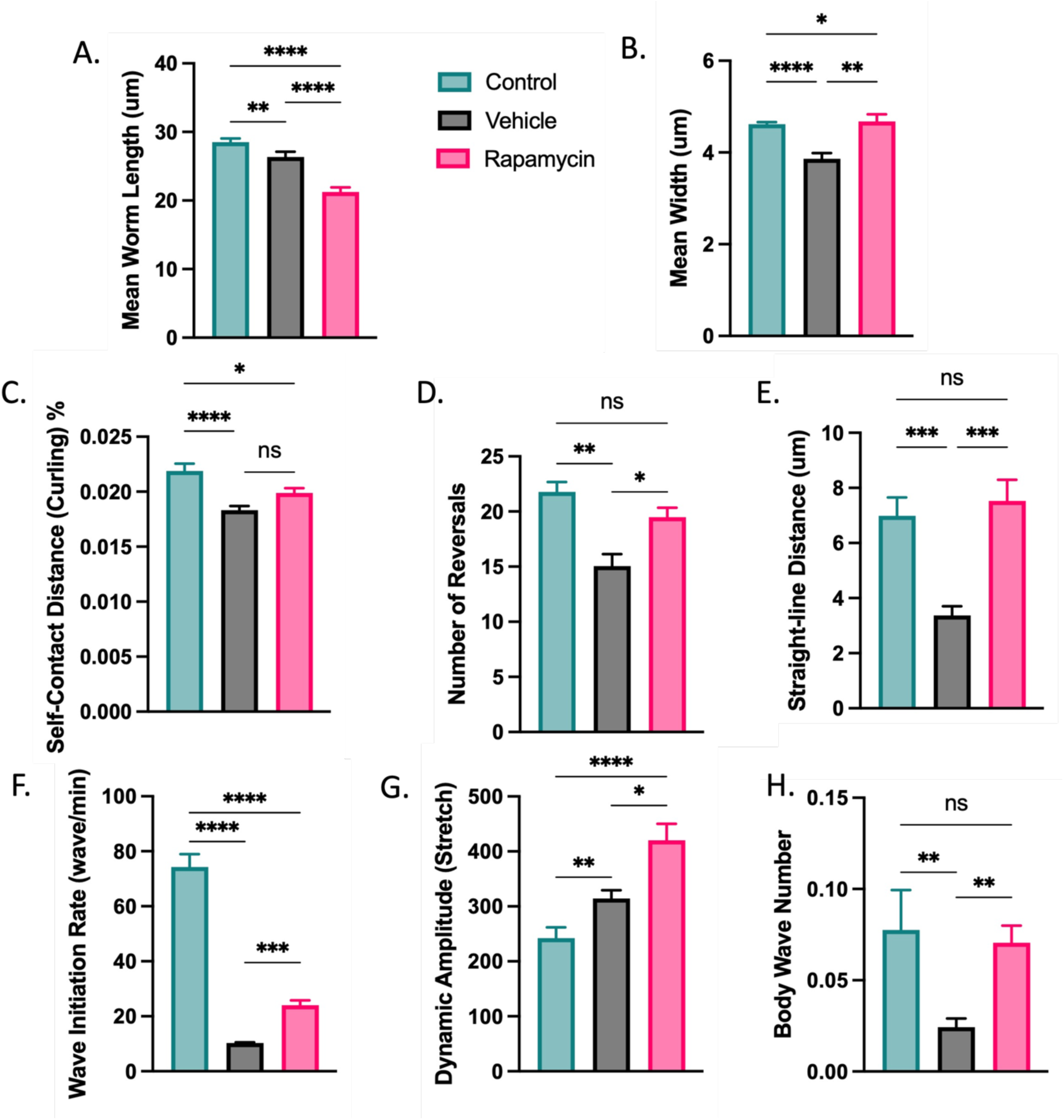
Rapamycin treatment alters body morphology and improves locomotor behavior in CL4176 strain. Quantitative analysis of WormLab (MBF Bioscience)-tracked morphology and locomotion in age-synchronized CL4176 worms cultured NGM containing either 0.2 % DMSO (Vehicle) or 20 μM Rapamycin. **(A)** Mean worm length. Rapamycin treatment reduced worm body length relative to both control and vehicle groups. **(B)** Mean body width. Vehicle treatment reduced width, whereas rapamycin-treated worms were wider than both vehicle-treated worms and controls. **Crawling Analysis: (C)** Self-contact distance (curling). Vehicle reduced curling relative to control but rapamycin-treated worms did not differ from vehicle-treated worms but remained significantly different from control stain. **(D)** Number of reversals. Vehicle reduced reversal frequency; rapamycin restored it to control levels. **(E)** Straight-line distance traveled. Vehicle reduced straight-line distance; rapamycin restored it to control levels. **Swimming analysis: (F)** Wave initiation rate (waves/min). Vehicle markedly reduced wave initiation rate: rapamycin improved the wave initiation rate but remains less than control strain waving rate. **(G)** Dynamic amplitude (stretch). Vehicle increased dynamic amplitude relative to controls, and rapamycin increased further. **(H)** Body wave number. Vehicle reduced body wave number; rapamycin restored it to control levels. Data are presented as mean ± SEM (n = ∼100 worms per group, from 3 independent experiments). Statistical significance was determined by Kruskal–Wallis test with Dunn’s multiple comparisons. ns, not significant; *p < 0.05; **p < 0.01; ***p < 0.001; ****p < 0.0001.

**Table 1.**
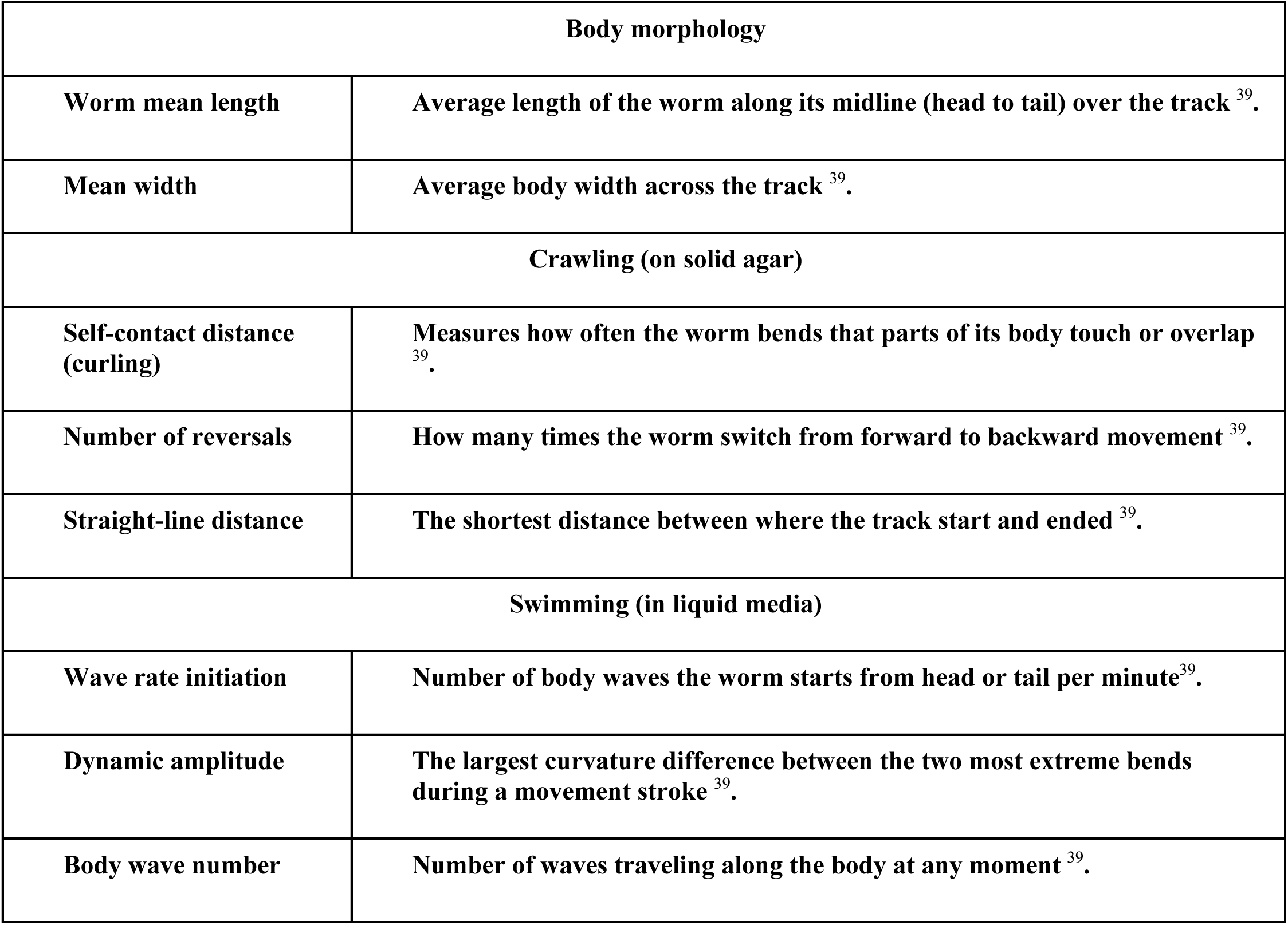
Definition of WormLab morphology and locomotion parameters.

### Rapamycin rescues the chemotaxis deficit in Aβ-expressing CL2355

Pan-neuronal Aβ_1–42_ expression in CL2355 produced the expected impairment in chemotaxis toward the volatile attractant benzaldehyde, with vehicle-treated worms displaying a mean chemotaxis index (CI) near zero. Rapamycin treatment significantly improved the CI in CL2355 in comparison to vehicle (one-way ANOVA, p = 0.008), restoring chemotactic behavior toward the level of non-Aβ transgenic controls (Figure 3). In contrast, rapamycin produced no significant change in CI in the non-transgenic control strain CL2122, whose chemotaxis was intact under both conditions.

**Fig 3.**
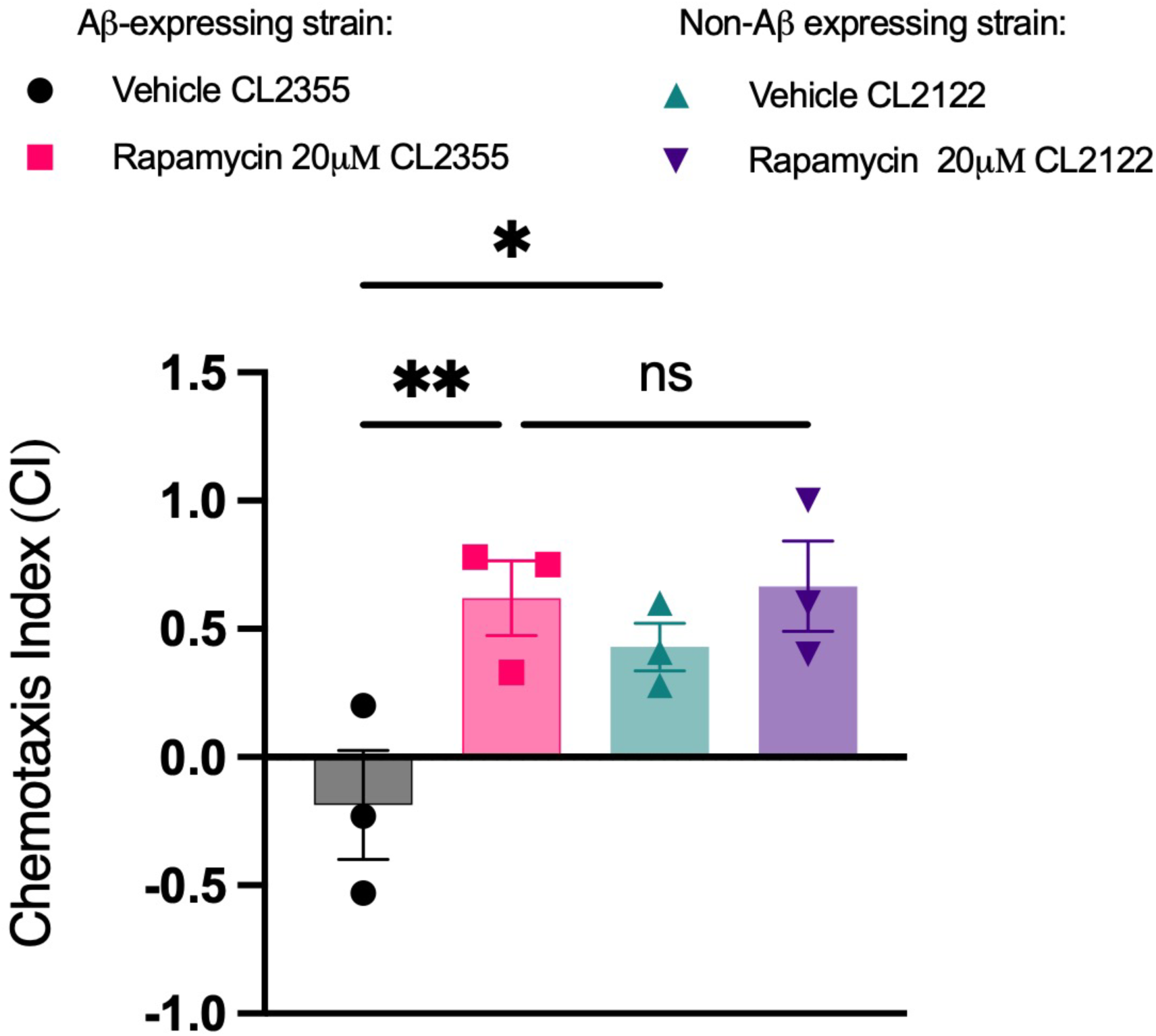
Rapamycin rescues the chemotaxis deficit in Aβ-expressing CL2355 without affecting the non-transgenic CL2122 control. Chemotaxis index (CI) toward benzaldehyde in CL2355 (Aβ_1–42_-expressing) and CL2122 (non-transgenic) worms treated with vehicle or 20 µM rapamycin. Vehicle-treated CL2355 worms showed an impaired CI near zero; rapamycin significantly increased CI in CL2355 (one-way ANOVA, p = 0.015), restoring chemotaxis toward control levels, with no significant effect in CL2122 (ns). Data are presented as mean ± SEM (n = ∼65 worms per group, from 3 independent experiments).

### Rapamycin extends lifespan and preserves healthspan in Aβ-expressing *C. elegans*

Kaplan– Meier survival analysis showed that vehicle-treated CL2355 worms underwent accelerated mortality relative to the non-Aβ transgenic CL2122 control, consistent with the toxic effect of pan-neuronal Aβ expression. Rapamycin significantly extended survival in CL2355 (log-rank, Bonferroni-adjusted p < 2 × 10⁻¹⁶), delaying complete population lethality across replicate experiments (Figure 4A), while producing at most a modest effect on CL2122 survival (Table 2). A parallel combined survival–locomotion assay, we show the overall activity as a proxy for the worms healthspan (Figure 4B). Activity was highest on day 1 and fell by day 3 in all groups. There was a small but significant main effect of treatment (F(1, 386) = 4.38, p = 0.037). The effect of rapamycin depended on strain, as shown by a significant treatment × strain interaction (F(1, 386) = 5.30, p = 0.022). In CL2355, rapamycin-treated worms tended to be more active than vehicle-treated worms, especially on days 3 and 5. In CL2122 controls, rapamycin tended to lower activity at early time points, most clearly on day 1. The three-way time × treatment × strain interaction was also significant (F(5, 386) = 3.92, p = 0.0018) (Table 3).

**Fig 4.**
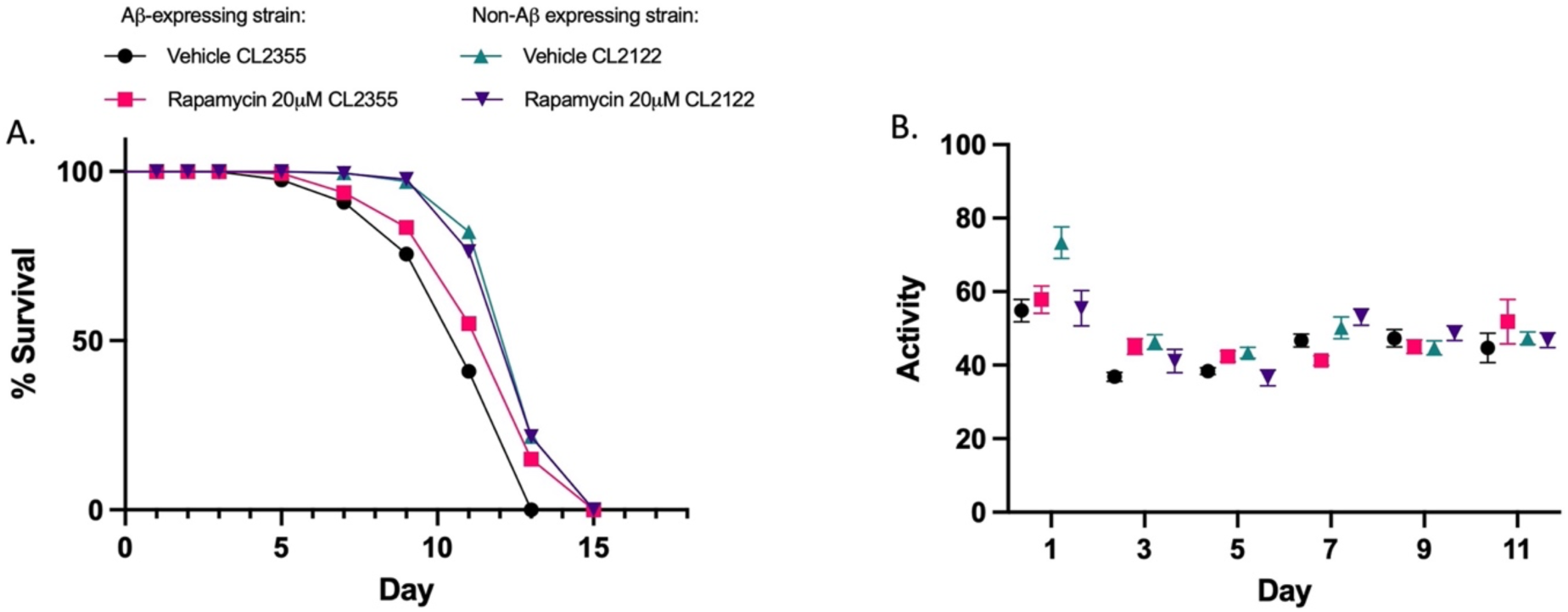
Rapamycin extends survival of Aβ-expressing transgenic strain and Improve overall Healthspan. Rapamycin extends survival in Aβ1-42-expressing transgenic *C. elegans*, with no effect in the control strain. Kaplan–Meier survival curves of transgenic CL2355 worms (pan-neuronal human Aβ1-42 expression) and control CL2122 worms, treated with vehicle (0.2% DMSO) or 20 µM rapamycin beginning at day 1 of adulthood and scored every other day. Rapamycin treatment increased survival relative to vehicle in the transgenic CL2355 strain but did not extend survival in the control CL2122 strain. Data represent n = 3 independent biological replicates (total worms per group ∼ 300). Statistical significance was determined by log-rank (Mantel–Cox) test with Bonferroni correction for multiple comparisons; all pairwise comparisons among the four groups were significant (p < 0.0001) multiple comparisons in Table1. **(B)** Activity was measured on days 1, 3, 5, 7, 9 and 11 in Vehicle CL2355 (black circles), Rapamycin CL2355 (pink squares), Vehicle CL2122 (teal triangles), and Rapamycin CL2122 (purple inverted triangles), using WormLab (MBF Bioscience). Activity was highest on day 1 across all groups; activity had declined in all groups and then remained relatively stable through day 11. Data are presented as mean ± SEM (n = ∼30 worms per group per day, from 2 independent experiments). Data were analyzed by three-way ANOVA with factor 1-treatment, factor 2-Strain and day as row factor. The treatment × strain interaction was significant (F(1, 386) = 5.30, p = 0.022). See Table 2.

**Table 2.** Pairwise comparisons of vehicle and rapamycin treatment across transgenic strain CL2355 and its control CL2122.

|  | **Vehicle CL2355** | **Vehicle CL2122** | **Rapamycin CL2355** |
|---|---|---|---|
| **Vehicle CL2122** | **<0.0001** | **-** | **-** |
| **Rapamycin CL2355** | **<0.0001** | **<0.0001** | **-** |
| **Rapamycin CL2122** | **<0.0001** | **0.78** | **<0.0001** |

**Table 3.** Three-way ANOVA of activity by time, treatment and strain.

| **Source of variation** | **F (DFn, DFd)** | **P value** | **% of total variation** |
|---|---|---|---|
| **Time** | **F (5, 386) = 28.73** | **< 0.0001 \*\*\*\*** | **23.73** |
| **Treatment** | **F (1, 386) = 4.381** | **0.0370 \*** | **0.72** |
| **Strain** | **F (1, 386) = 0.2815** | **0.5960 ns** | **0.05** |
| **Time × treatment** | **F (5, 386) = 1.757** | **0.1208 ns** | **1.45** |
| **Time × strain** | **F (5, 386) = 1.621** | **0.1533 ns** | **1.34** |
| **Treatment × strain** | **F (1, 386) = 5.297** | **0.0219 \*** | **0.87** |
| **Time × treatment × strain** | **F (5, 386) = 3.922** | **0.0018 \*\*** | **3.24** |

### Rapamycin dose not lower Aβ species levels

To test whether rapamycin affects Aβ levels, we ran western blots on worm lysates with anti-Aβ antibodies and resolved four immunoreactive species at approximately 4 (monomer), 20, 25, and 60 kDa (oligomers) from the muscle and neuronal Aβ strains treated with vehicle or 20 µM rapamycin. Rapamycin-treated worms showed a visual weaker signal for both oligomers and monomers (Figure. 5A). In rapamycin-treated muscle-strain worms, the 20 kDa species was about 45% lower than in vehicle-treated worms (Figure. 5B), the 16 kDa species approximately 48% lower (Figure. 5C), the 10 kDa species approximately 39% lower (Figure. 5D), and the ∼4 kDa monomer is approximately 17% lower (Figure. 5E). None of these differences were statistically significant (unpaired t-test, p > 0.05). In the neuronal strain, the Aβ signal was much weaker overall, and the most visible band was at about 16 kDa. Rapamycin lowered the 16 kDa and 20 kDa species by about 54% and 52% (Figure 5F-G), but no significant difference was detected.

**Figure 5.**
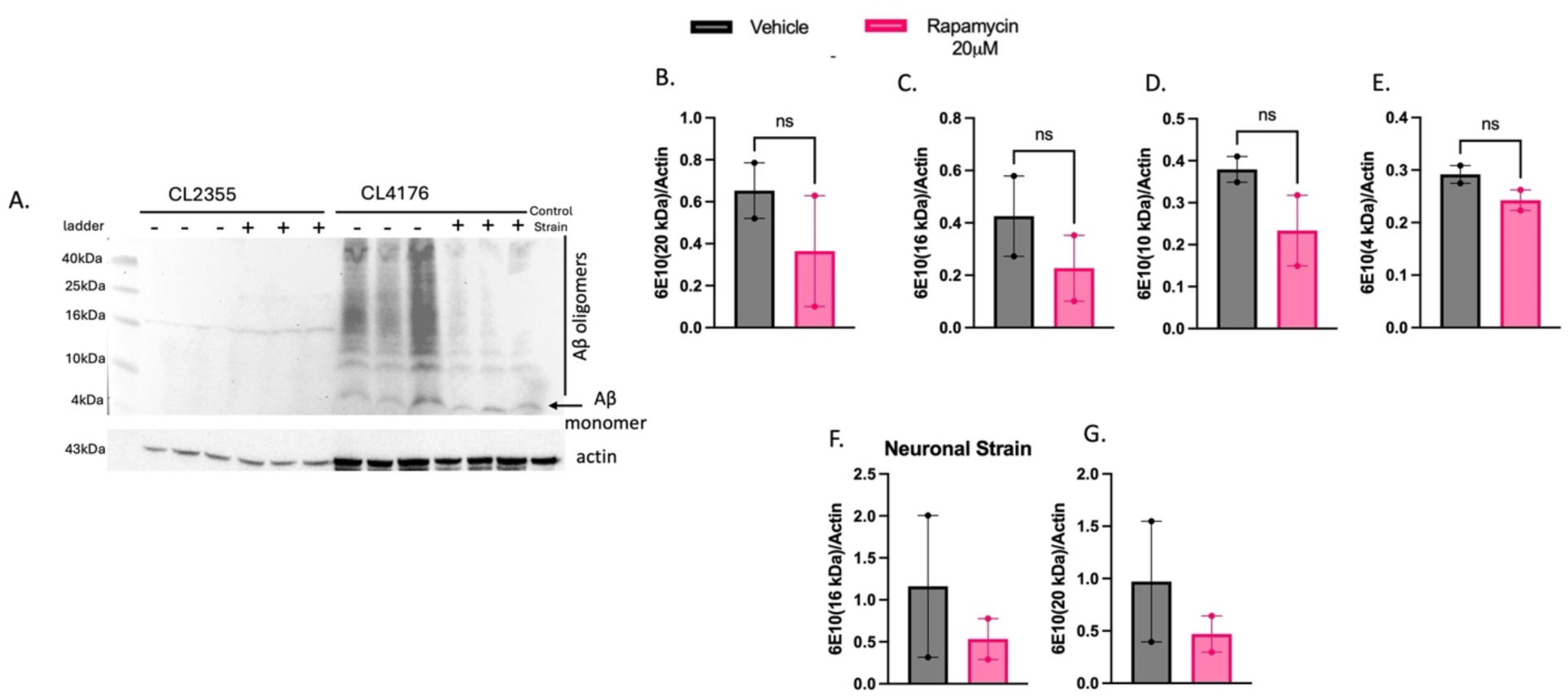
Rapamycin does not affect Aβ levels in neuronal or muscle Aβ-expressing *C. elegans* strains. **(A)** Representative western blot of Aβ species detected with the 6E10 antibody in lysates from the neuronal Aβ strain (CL2355) and the muscle Aβ strain (CL4176) treated with vehicle DMSO, 0.2% (-) or 20 µM rapamycin (+). The control strain (CL2122]) was included as a negative control. Aβ oligomers (∼10–40 kDa) and Aβ monomer (∼4 kDa) are indicated. Actin (43 kDa) was used as a loading control. Quantification of 6E10-immunoreactive bands at (B) 20 kDa, (C) 16 kDa, (D) 10 kDa and (E) 4 kDa in the muscle strain, normalized to β-actin all of them did not show any significance. **(F, G)** Quantification of 6E10-immunoreactive bands at (F) 16 kDa and (G) 20 kDa in the neuronal strain, normalized to actin with. Data are presented as mean ± SEM, with individual points representing 2 independent biological replicates (average of 3 lanes per sample). Statistical significance was determined by unpaired two-tailed t-test.

## Discussion

This study provides a systematic, vehicle-controlled assessment of rapamycin’s neuroprotective effects across muscle-and neuron-expressing *C. elegans* models of Aβ proteotoxicity. The delay in Aβ-induced paralysis, improvement in the number of reversal and straight-line distance along with swimming parameters suggest that rapamycin restores neuromuscular function in body-wall muscle. Because crawling and swimming are controlled by partially distinct interneuron populations, the consistency of the effect across both behaviors indicates a systemic improvement in the proteostatic environment of Aβ-expressing muscle, consistent with prior reports that mTORC1 inhibition enhances the clearance of Aβ ^9,27^.

Rapamycin rescued the chemotaxis deficit in CL2355, without any measurable effect on the non-Aβ transgenic CL2122 control. Chemotaxis toward benzaldehyde depends on intact Amphid Wing C cells (AWC) / Amphid Wing A cells (AWA) olfactory neuron signaling ^22,28^. The strain-selective nature of the rescue which is present in the Aβ-expressing strain but absent in the transgenic control suggests that rapamycin provides neuroprotection and alleviates Aβ-driven sensory impairments rather than exerting a generalized effect on sensation or motor output. This pattern parallels the selectivity of rapamycin’s cognitive benefits in rodent AD models ^11^, where improvements are observed in transgenic but not the transgenic control worms, suggesting that CL2355 captures a functionally analogous form of Aβ-specific dysfunction.

In the context of lifespan, some interventions protect Aβ-expressing *C. elegans* by slowing aging generally: apigenin, for example, extends lifespan and healthspan in both wild-type and Aβ-expressing worms^29^. In contrast, the pronounced extension of Kaplan–Meier survival in rapamycin-treated CL2355 and the absence of effect on the non-Aβ transgenic CL2122 control suggests that its benefit reflects on mitigation of Aβ-specific toxicity rather than generic longevity extension (by its well-established pro-longevity effects on the mTOR pathway) ^30,31^.Importantly, the combined survival–locomotion analysis demonstrated that rapamycin improves healthspan rather than simply prolonging survival in an impaired state ^32^. The progressive delay in the onset of motor decline in rapamycin-treated CL2355 worms, indicates that the drug maintains neuromuscular integrity over time. In aging research, the distinction between lifespan extension and healthspan improvement is of great importance: an intervention that extends life without preserving function is of limited therapeutic value ^33,34^. The present data demonstrate that rapamycin achieves both in the Aβ proteotoxic context.

Rapamycin treatment improved behavioral outcomes and healthspan alongside lifespan extension in CL4176 and CL2355 worms without detectable reduction in the Aβ monomer or oligomer burden. Two potential mechanisms may be accounted for the unmeasurable difference. First, rapamycin alters Aβ compartmentalization through modulation of the autophagy–lysosome pathway, although protective sequestration within autophagic vesicles is not addressed in the present study. Rapamycin has been shown to restore HLH-30 nuclear localization in an Aβ-expressing *C. elegans* background ^35^. Secondly, chaperone-associated sequestration has been linked to reduced Aβ toxicity without a reduction in overall peptide accumulation, providing a platform for protection through altered Aβ distribution rather than reduced abundance ^36^.

In this study, rapamycin improved behavioral performance in muscle-and neuron-expressing *C. elegans* models of Aβ proteotoxicity. Lifespan extension was observed alongside delayed locomotor decline. These benefits occurred without a detectable reduction in Aβ monomer or oligomer levels, indicating that functional improvement was not accompanied by a measurable decrease in the Aβ species burden. Altered Aβ compartmentalization and enhanced cellular resistance to proteotoxic stress is possible but unconfirmed hypothesis. As a future direction, we recommend examining Aβ subcellular localization^37^ and autophagic flux^38^ and assess the contribution of protective stress-response pathways^38^.

The pace of conventional drug discovery: from target identification and de novo compound synthesis through Phase III trials, routinely spans fifteen to twenty years and costs well over a billion dollars per approved agent resulting in limited disease-modifying options ^2^. The traditional pipeline timeline poorly matches to a disease whose prevalence is projected to more than double within the next three decades ^2^.Drug repurposing is redirecting compounds with established human safety profiles, known pharmacokinetics, and existing regulatory approval toward new therapeutic indications will offer a strategically compelling alternative that compresses development timelines, reduces cost, and substantially de-risks early-phase clinical investigation ^12^.Progress in AD drug discovery remains constrained by the cost and throughput limitations of mammalian models ^13^. These findings will support further investigation of rapamycin as a modulator of Aβ-associated proteotoxicity and highlight the importance of integrating functional outcomes with biochemical measures of autophagy markers.

